# Sequence-Derived Geometric Priors Enable Controllable Generation of Intrinsically Disordered Protein Ensembles

**DOI:** 10.64898/2026.11.19.768000

**Authors:** Wonkyeong Jang, Woong-Hee Shin

## Abstract

Predicting heterogeneous conformational ensembles from sequence remains a central challenge for intrinsically disordered proteins (IDPs). Sequence-to-ensemble models often entangle geometric inference with coordinate generation, obscuring error sources and limiting control over ensemble properties. Here we introduce MANTA, a Cα-level framework that separates these stages. An encoder derives geometric priors from protein language-model representations, and a graph-Laplacian decoder converts these priors into conformational ensembles. On a homology-held-out test set, replacing predicted priors with reference-derived priors under the same decoder reduces the median absolute relative radius-of-gyration error from 8.78% to 1.89%, identifying geometric-prior estimation as the dominant bottleneck in MANTA. Across an external benchmark of 27 IDPs, MANTA generates broad, locally valid ensembles. Decoder controls modulate mean compactness and conformational breadth along partially separable axes, while compactness modulation alters motif exposure in tau and amyloid-β. Meanwhile, α-synuclein variants shift local exposure populations at nearly unchanged global size. MANTA provides a diagnostic and controllable framework for ensemble generation that captures both global dimensions and local conformational organization.

## 1. Introduction

Intrinsically disordered proteins (IDPs) and regions (IDRs) cannot adopt a single stable folded structure ^[1]^, yet mediate diverse biological functions through heterogeneous conformational ensembles^[2]^. Around 70% of human proteins contain at least one IDR of 30 residues or longer^[2]^. As there is no single native state, their behavior is governed by ensemble-level properties, including chain compactness, long-range conformational preferences, and transient intermolecular contacts^[3]^. IDP/IDR modeling therefore diverges from conventional protein structure prediction: the goal is not to recover a single native state but to generate a geometrically plausible distribution of conformations.

Existing approaches address the problem from contrasting directions. Atomistic and coarse-grained (CG) simulations generate ensembles from physics-based rules^[4,5]^, but their validity depends on parameterization, sampling, and the chosen physical models^[4,6]^. Conversely, polymer-based and statistical models capture global dimensions such as radius of gyration (Rg) and end-to-end distance, and sequence-dependent trends, but rely on prescribed interaction models, scaling assumptions, or calibration to experimental observables^[5,7]^.

Deep-learning methods, including coordinate-generative and diffusion-based models, have recently advanced ensemble generation^[6,8-10]^. Sequence-driven generative methods fill a distinct niche — probing how much ensemble-level geometry can be inferred from sequence and realized rapidly at scale — but in many cases the transition from sequence to ensemble properties remains implicit, making errors in compactness, distance geometry, or contact organization hard to trace to prior inference, coordinate realization, or limited control over heterogeneity^[6,8-10]^.

Protein language models (PLMs) provide a natural source of sequence-derived information by encoding biochemical and evolutionary patterns directly from amino-acid sequences^[11]^. However, PLM representations do not themselves define conformational ensembles^[2,3]^, and for disordered proteins, predicted distances or contacts are better interpreted as statistical geometric tendencies than as deterministic structural constraints. The key challenge is therefore not only to infer such tendencies from sequence, but also to determine which aspects of ensemble organization they reliably encode and how they can be translated into heterogeneous conformational populations.

In this study, three research questions are posed: the extent to which ensemble-level geometry can be inferred from sequence-derived representations; whether errors in sequence-to-ensemble arise primarily from geometric inference or from coordinate realization; and whether separating these processes enables local conformational changes to be resolved beyond global measures such as Rg. To address these questions, we developed MANTA (Modular Architecture for Navigable Topological Assembly), a sequence-based Cα-level framework for IDP ensemble generation that separates geometric-prior estimation from conformational realization. The encoder of the system converts PLM-derived representations into several different signals, including expected inter-residue distances, confidence-weighted pair constraints, local chain geometry, and compactness tendencies. A graph-Laplacian decoder then realizes these priors as heterogeneous ensembles, with controllable mean-size and breadth settings. The modular design of the system enables the evaluation of sequence-derived information and coordinate generation to be conducted independently, while concurrently providing direct control over ensemble compactness and conformational heterogeneity.

## 2. Results

### 2.1. Overview of MANTA

The IDP ensemble generation incorporates two components — estimating sequence-dependent geometric tendencies and realizing them as heterogeneous conformational populations — that are often merged into a single prediction problem. However, MANTA addresses them separately. The encoder converts fixed protein-language-model features into sequence-derived geometric priors: expected Cα–Cα distances; pair-confidence estimates that indicate the relative reliability of each predicted pairwise distance and modulate its weight during decoding; local chain-geometry terms that describe *i, i+2* Cα–Cα distances constraining local chain organization; and a compactness-related output, an Rg estimate derived from the predicted distance map. The decoder generates Cα-level ensembles from the encoder outputs through frame-specific Rg sampling, Rg-scaled graph construction, Laplacian-guided SMACOF (scaling by majorizing a complicated function) relaxation^[12]^ — an iterative multidimensional-scaling procedure that updates three-dimensional coordinates to minimize the weighted discrepancy between prescribed and realized distances — followed by local scaling, and final global Rg matching **(Figure 1)**.

**Figure 1.**
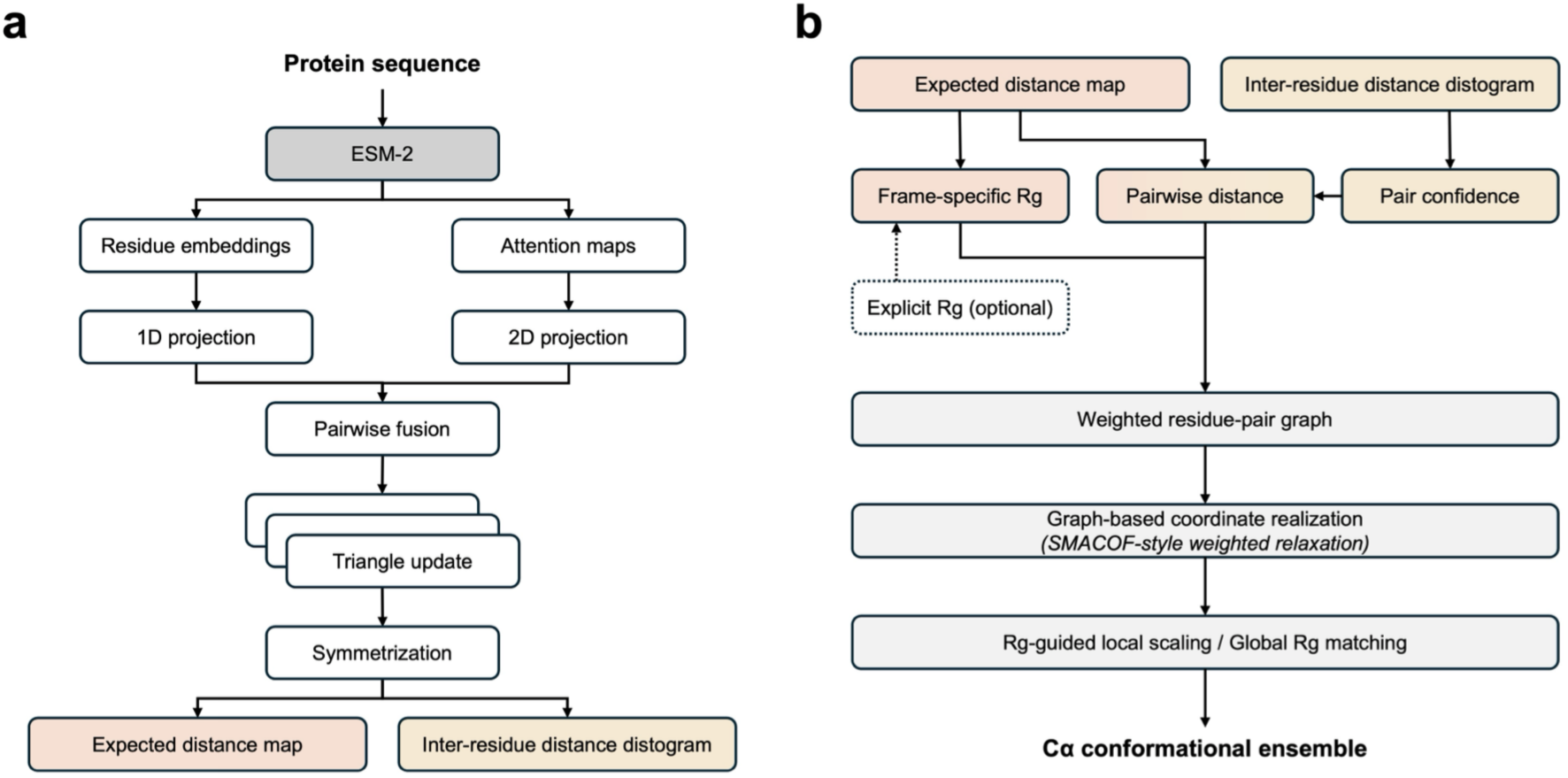
Overview of the MANTA encoder–decoder framework. **a.** The encoder converts fixed protein-language-model features into sequence-derived geometric priors as outputs, including an expected distance map and an inter-residue distance distogram. **b.** The decoder samples frame-specific Rg and pairwise distance targets, derives pair confidence from the distogram, constructs a weighted residue-pair graph, and generates Cα ensembles through graph-based SMACOF relaxation, followed by Rg-guided local scaling and global Rg matching, with an explicit Rg optionally supplied for target-conditioned generation.

MANTA supports three different operating modes. In the default sequence-only mode, compactness and distance geometry are inferred from the given IDP sequence alone. In the Rg-scale-controlled mode, sequence-derived priors are decoded under shifted mean-size settings without retraining the encoder. In the explicit Rg-conditioned mode, a user-supplied target compactness is passed directly to the decoder. MANTA is therefore not only a sequence-to-ensemble generator but a framework that separates geometric-prior estimation, coordinate realization, mean compactness control, and ensemble-width control.

### 2.2. Capturing Geometric Features of IDP from Sequence with the Encoder

First, the encoder was assessed for its ability to reproduce experimentally observed geometric features on a homology-held-out test set of 675 ensembles from the Protein Ensemble Database (PED)^[13]^ using MMseqs2^[14]^-based clustering at 30% sequence identity and 0.5 coverage. The Rg inferred from the predicted pairwise distance map correlated strongly with the reference ensemble Rg, with a Pearson’s correlation coefficient (PCC) of 0.914 and a mean absolute error (MAE) of 4.00 Å; a global multiplicative calibration factor of 1.1102, optimized on the validation set with Optuna^[15]^, reduced the MAE to 3.84 Å at the same PCC **(Figure 2a,b)**. The MANTA encoder also demonstrated the capacity to predict pairwise Cα distance maps (Figure 2c). Predicted pairwise Cα distance maps showed a median all-pair MAE of 6.45 Å and a median nonlocal MAE (pairs separated by at least three residues) of 6.41 Å, with a median per-target PCC of 0.938 against reference maps (Figure 2c). The corresponding median root-mean-square error (RMSE) values were 8.26 Å and 8.28 Å, respectively. This error depended on the sequence separation: short-(3-5) and medium-range (6-20) residue pairs were more accurate, while long-range pairs (> 20) exhibited larger, more variable errors (Figure 2d). Sequence-derived priors thus contain substantial short-and medium-range geometric information, but long-range IDP geometry remains uncertain.

**Figure 2.**
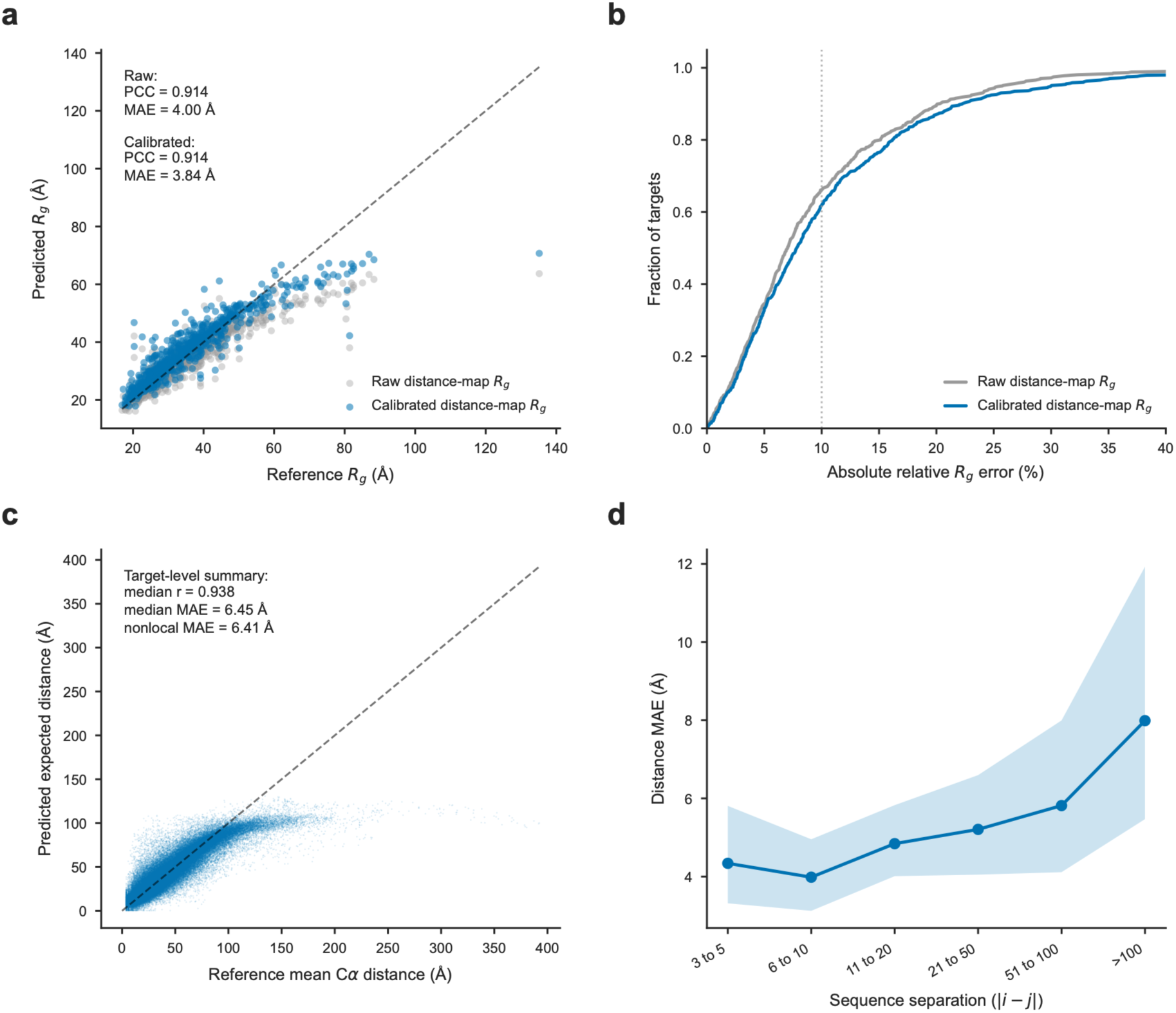
**a.** Reference versus predicted Rg before and after scalar calibration. **b.** Empirical cumulative distribution function of absolute relative Rg error. The dotted line marks 10% error. **c.** Predicted expected Cα distances compared with reference ensemble-averaged distances. **d.** Distance MAE by sequence separation; points show medians and shading shows the interquartile range.

### 2.3. Analyzing the Source of Error by Decoupling the Encoder and Structure Generation

The modularity of MANTA allows the residual error of generated ensembles to be attributed either to imperfect encoder priors or to the coordinate-generation step. An ensemble of 300 conformers was generated under two fixed settings with identical decoder hyperparameters: encoder-predicted priors and reference-derived priors computed from the reference ensemble structures. With predicted priors, the decoder reproduced global compactness (median signed relative Rg error of 5.90%, a median absolute relative Rg error of 8.78%) and locally valid geometry, with a median all-pair distance MAE of 8.96 Å, a median Jensen–Shannon (JS) divergence of 0.010 between generated and reference pairwise distributions, and a contact-density PCC of 0.722 **(Supplementary Table S1)**.

Supplying reference-derived priors improved every metric: median absolute Rg error fell from 8.78% to 1.89%, all-pair and nonlocal distance MAE to 3.89 Å and 3.96 Å (a 56-57% reduction), JS divergence to 0.004, and contact-density PCC increased from 0.722 to 0.848 (Supplementary Table S1, **Figure 3**). Therefore, the graph-Laplacian decoder reliably generates IDP-like ensemble distributions when accurate geometric information is given. The primary residual error of sequence-only generation is prior estimation — particularly long-range geometry — rather than coordinate realization.

**Figure 3.**
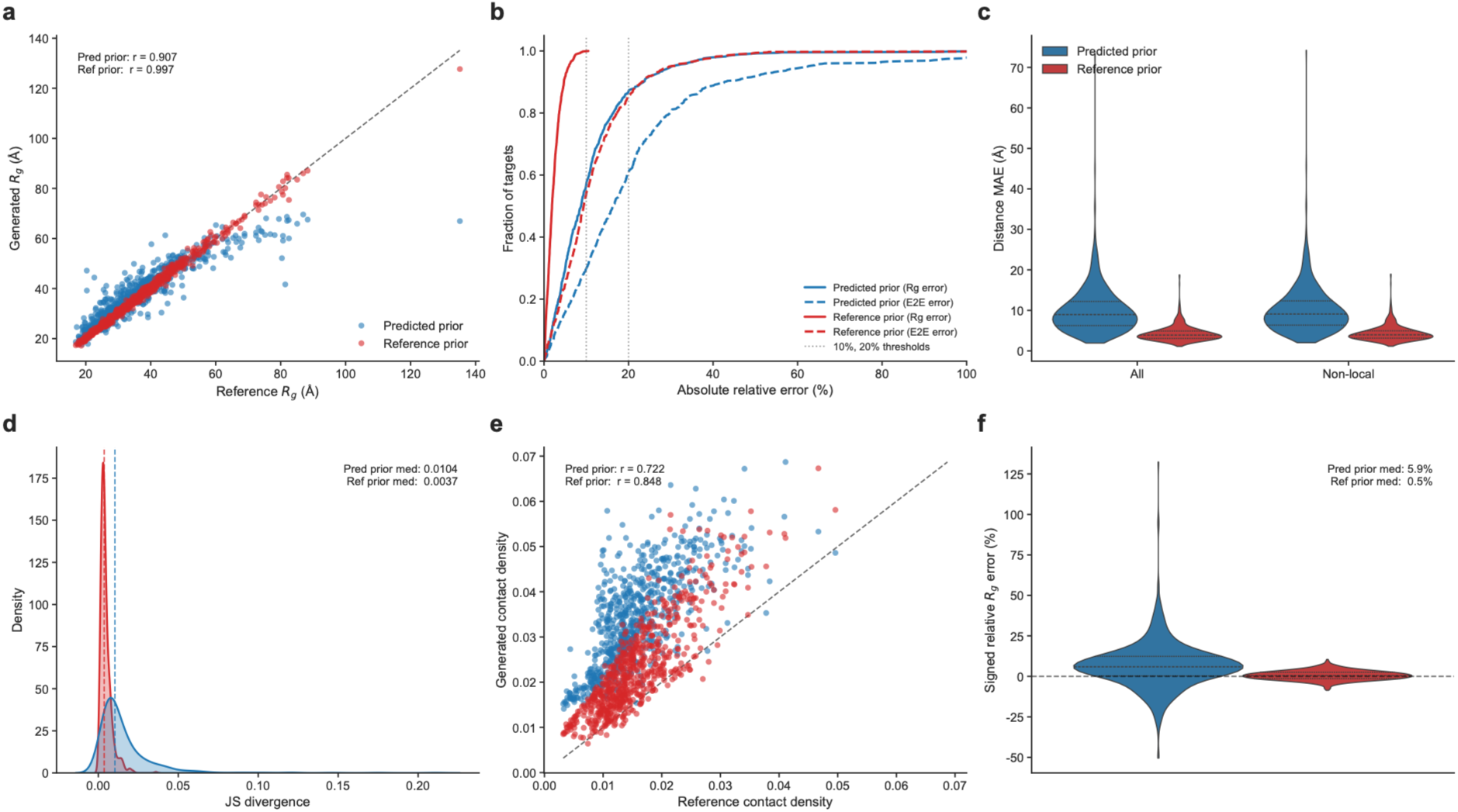
Comparison of MANTA performance between predicted prior and reference prior. **a.** Reference versus generated ensemble-averaged Rg for ensembles generated from encoder-predicted priors and reference-derived priors. **b.** Empirical cumulative distributions of absolute relative Rg and end-to-end distance errors. Vertical dotted lines indicate 10% and 20% error. **c.** Distributions of all-pair and nonlocal pairwise distance MAE. **d.** JS divergence between reference and generated pairwise distance distributions. Dashed lines indicate median values. **e.** Reference versus generated contact density. **f.** Distributions of signed relative Rg error. The same decoder hyperparameters were used for both prior sources.

### 2.4. The Frozen Sequence-Only Mode Defines the External Benchmark Operating Point

We next benchmarked the sequence-only mode on the IDPFold^[9]^ test set of 27 IDPs, with the decoder Rg-scale proxy *p_Rg_* fixed at 1.00. All 27 systems were excluded from training, validation, Rg scaling, and hyperparameter and *p_Rg_* selection to prevent leakage. The frozen default produced a mean signed fractional Rg error (ε_Rg_), defined as the generated minus experimental Rg, divided by the experimental reference Rg, of −0.11, indicating slightly compact ensembles relative to experiment. This is the default operating point, not an Rg-optimized state; because MANTA operates at Cα-level, cross-method comparison was restricted to the resolution-compatible signed ε_Rg_, for which MANTA was comparable to other IDP ensemble generators **(Supplementary Table S2)**.

To confirm that generated ensembles represented continuous conformational populations rather than isolated compactness-matched conformers, the MANTA and CALVADOS2^[5]^ ensembles were projected onto Rg-RMSD (root-mean-square-distance) occupancy maps for all 27 systems **(Supplementary Figure S1)**. MANTA occupied broad, continuous regions with distributions and coverage similar to the MD-based CALVADOS2 across most systems, despite generating ensembles through an encoder-decoder framework rather than simulation. Local geometry was valid in 100% of benchmark conformers, confirming that external compactness evaluation did not induce local Cα-geometry violations. The full sequence-to-ensemble pipeline ran in a median of 17.14 s (mean 21.02 s) per system on an NVIDIA RTX 2080 Ti, with feature extraction and encoder inference contributing a negligible fraction and the decoder and post-processing dominating **(Supplementary Table S3)**. MANTA thus offers rapid Cα-level sequence-to-ensemble generation.

### 2.5. Decoder Controls Separate Mean Compactness from Conformational Breadth

To investigate whether the negative ε_Rg_, observed in the default mode, reflected a fixed sequence-derived bias or a shiftable decoder operating point, the decoder Rg-scale proxy, *p_Rg_*, was varied at constant frame-level Rg sampling width; *p_Rg_* below one imposes compacting settings, one is the frozen default, and above one imposes expanding settings. Increasing *p_Rg_* produced coordinated expansion across observables: median Rg and end-to-end distance increased monotonically. At the same time, contact density fell, with the within-ensemble Rg standard deviation changing only modestly — consistent with *p_Rg_* acting on mean ensemble size rather than breadth (**Figure 4a**).

**Figure 4.**
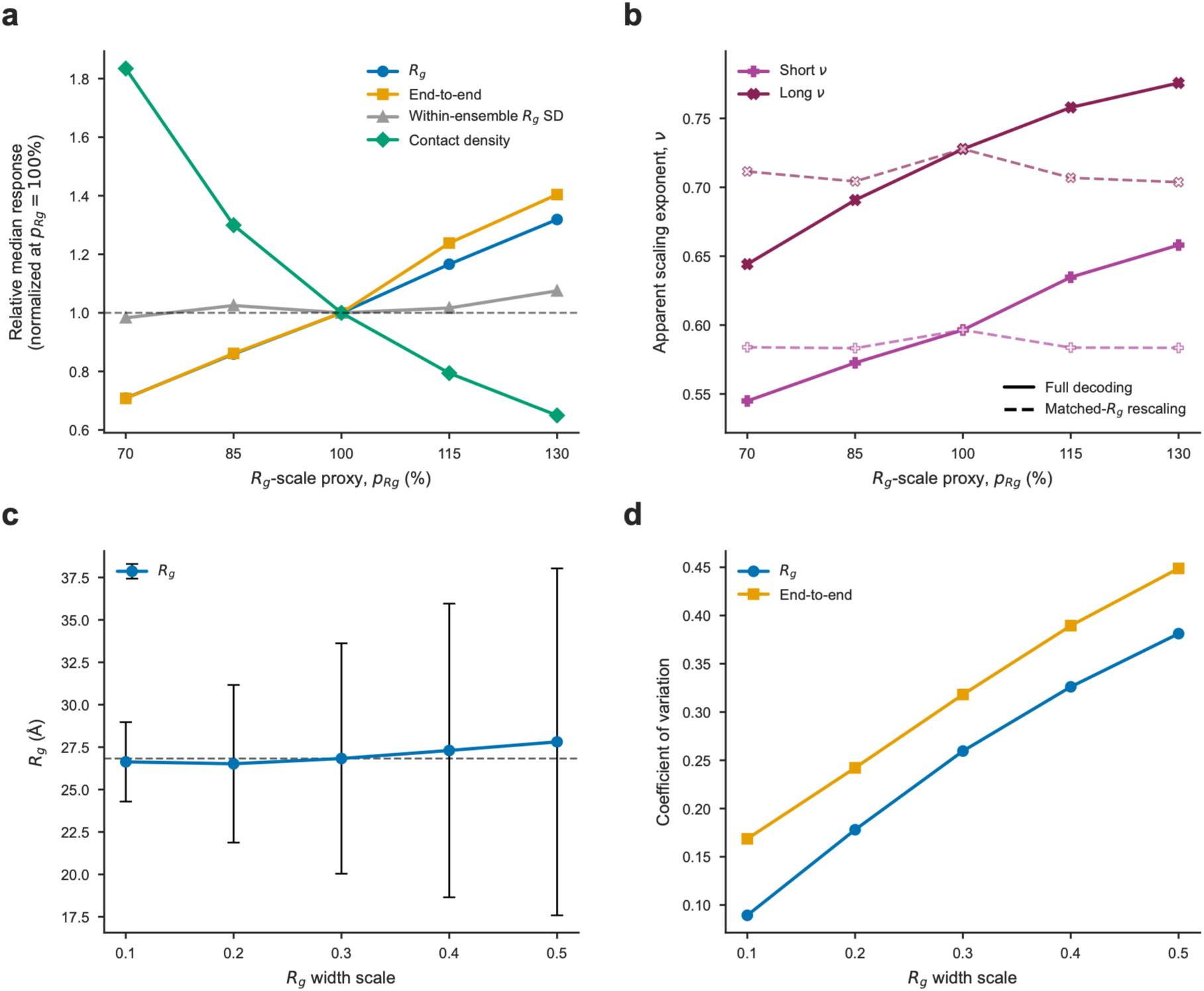
Mean compactness, apparent internal scaling and conformational heterogeneity modulated by decoder controls. **a.** Median ensemble observables across the decoder Rg-scale proxy, *p_Rg_*, normalized to the frozen default sequence-only setting (*p_Rg_*=1.00; 100%). The dashed horizontal line indicates the normalized response at *p_Rg_*=1.00. **b.** Apparent short-and long-regime sequence-separation scaling exponents under full Rg-scale-controlled decoding and matched-Rg final-rescale-only controls. **c.** Benchmark-wide median generated Rg across Rg width scales, with the Rg-scale proxy held fixed at the default setting: median of the ensemble-averaged Rg values, with whiskers showing the median of the within-ensemble Rg standard deviations and therefore representing conformational spread rather than uncertainty in the central estimate. The dashed line marks the median generated Rg at the default Rg width scale of 0.30. **d.** Benchmark-wide median coefficients of variation for Rg and end-to-end distance across Rg width scales.

The Rg-scale modulation also altered apparent internal chain scaling, measured by the sequence-separation scaling exponent ν, which describes how the spatial distance between residues scales with their distance along the chain. As *p_Rg_* increased from 0.70 to 1.30, the apparent short-range (1-10 residues) scaling exponent rose from 0.545 to 0.658, and the long-range exponent rose from 0.644 to 0.776 (Figure 4b). At default setting (*p_Rg_* = 1.00), the short-and long-range exponents were approximately 0.60 to 0.73, respectively. The long-range exponent exceeded the good-solvent excluded-volume reference value (ν ≈ 0.588^[16]^) across the full sweep. These exponents describe how mean Cα-Cα distance scale with sequence separation over the fitted ranges. To confirm this reflected decoder-level generation rather than trivial scaling, we built matched-Rg final-rescale-only controls by globally rescaling default conformers to the conformer-level Rg distributions of the Rg-scale-controlled ensembles. Both exponents increased monotonically under full decoding but stayed nearly flat in the controls, so the Rg-scale response reflects altered internal organization, not post hoc coordinate scaling. On the IDPFold benchmark, the same sweep shifted ε_Rg_ from -0.367 at *p_Rg_* = 0.70, through -0.236, -0.114, 0.023, to 0.153 at *p_Rg_* = 1.30 **(Supplementary Table S4)**, showing the default compactness bias is controllable without retraining the encoder or using external Rg values to redefine the default model.

Holding *p_Rg_* fixed and varying the Rg width scale broadened ensembles with only modest central-Rg change: median Rg changed from 26.63 Å at a width scale of 0.1 to 27.81 Å at 0.5, while the median Rg coefficient of variation increased from 0.089 to 0.381 and the median end-to-end coefficient of variation increased from 0.169 to 0.449 (Figure 4c,d). Mean compactness and conformational heterogeneity are therefore partially separable decoder variables. Finally, explicit Rg conditioning reproduced supplied targets almost exactly (mean signed ε_Rg_ 0.0014, median 0.008; mean absolute ε_Rg_ 0.0148), so MANTA can act as either a sequence-only generator or a compactness-guided decoder when target information is available.

### 2.6. Sequence Variation Reshapes Local Conformational Exposure Beyond Global Chain Dimensions

Rg captures global chain dimensions but does not describe how specific motifs are buried or exposed within an ensemble. For aggregation-related IDPs, such local accessibility is particularly relevant because intramolecular shielding can modulate the availability of aggregation-prone motifs for intermolecular interactions. We therefore examined tau, amyloid-β (Aβ), and α-synuclein using Cα contact-based exposure descriptors.

The tau repeat domain construct K18 contains four pseudo-repeats (R1-R4), whereas K19 lacks R2^[17]^. K18 therefore contains both aggregation-associated paired helical filaments hexapeptide (PHF6) motifs, PHF6* in R2 and PHF6 in R3, whereas K19 only PHF6^[18]^. Increasing *p_Rg_* raised contact-defined exposure and reduced the fraction of motif residues with nonlocal contacts for all three construct-motif combinations (**Figure 5a,b**). Across all tested settings, PHF6 was more exposed and less contacted in K19 than in K18. This difference is consistent with the absence of R2 in K19. These Cα-level monomeric ensembles quantify geometric accessibility. Assessing nucleation competence also requires information about side-chain packing^[19]^.

**Figure 5.**
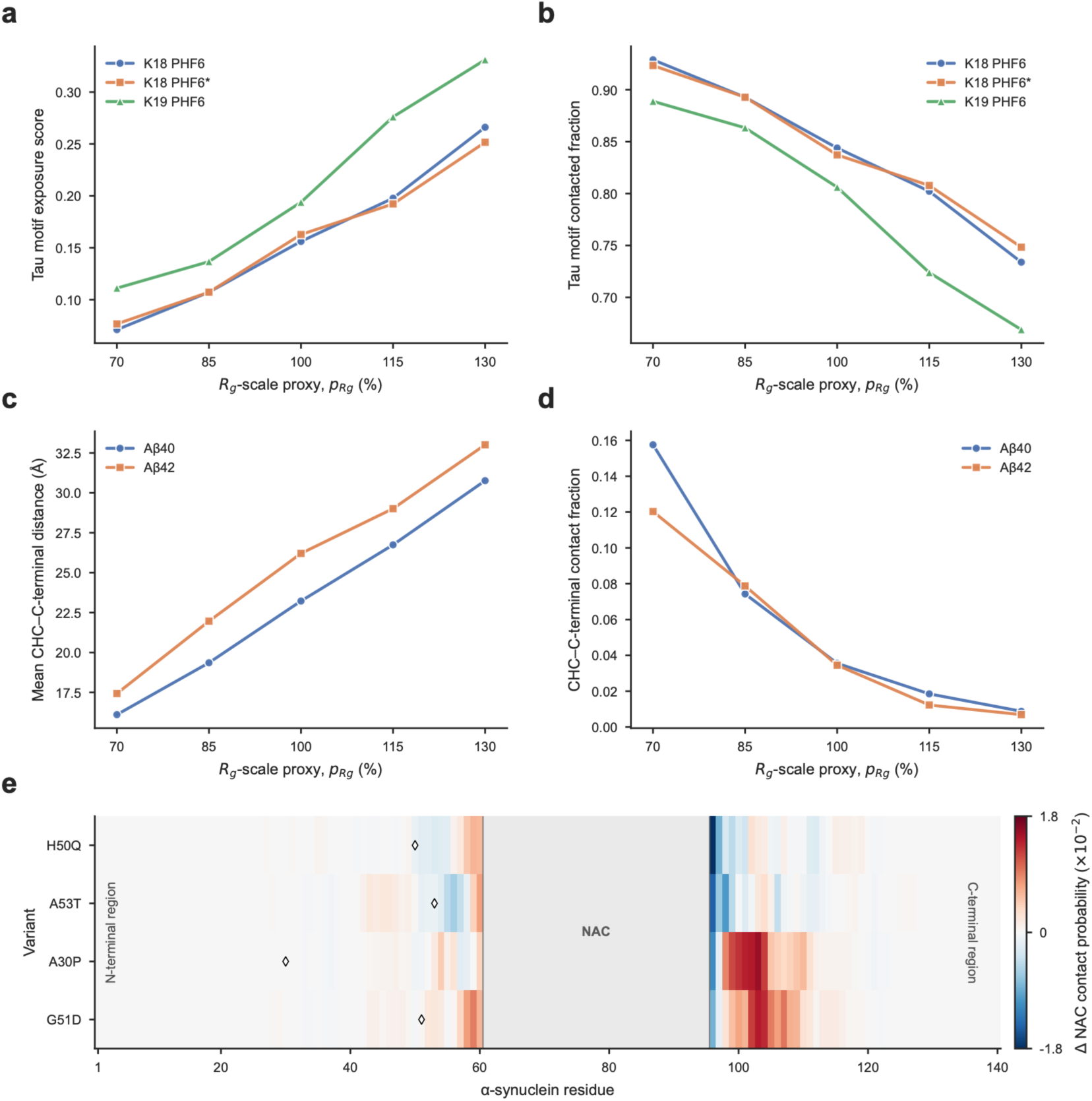
Rg-scale-controlled ensembles project onto regional exposure readouts. **a.** Contact-defined exposure of tau aggregation-associated motifs across Rg-scale settings. **b.** Contacted fraction of the tau PHF6 motifs. **c.** Mean Cα-level separation between the central hydrophobic cluster and C-terminal region in Aβ40 and Aβ42. Both peptides show increased CHC–C-terminal separation with Rg-scale-driven expansion, while Aβ42 maintains larger separation across the Rg-scale range. **d.** CHC–C-terminal contact fraction in Aβ40 and Aβ42. Rg-scale-driven expansion reduces CHC–C-terminal contact in both peptides. **e**. Residue-wise changes in NAC contacts of α-synuclein variants relative to WT. Diamonds indicate the mutation sites.

Aβ aggregation is driven by two hydrophobic regions: the central hydrophobic cluster (CHC, residues 17-21, LVFFA)^[20]^, which contains the F19/F20 aromatic pair that promotes β-sheet formation^[21]^, and the C-terminal region (31-42, IIGLMVGGVVIA)^[21]^, where I41 and A42 stabilize the β-strand and enhance hydrophobic packing^[22]^ relative to Aβ40. Interactions between these regions contribute to collapsed conformations and β-hairpin populations in Aβ monomers^[23]^. Increasing *p_Rg_* raised the mean CHC–C-terminal distance and lowered their contact fraction in both peptides (Figure 5c,d). Aβ42 showed greater CHC– C-terminal separation across the sweep. Contact fractions decreased for both peptides and were lower for Aβ42 at most settings. At Cα resolution, these differences reflect sequence-dependent geometric organization rather than the presence of a defined β-hairpin or differences in aggregation competence.

We next asked whether sequence variation could redistribute local conformational populations without substantially altering global chain dimensions. α-synuclein contains the aggregation-critical non-amyloid-β component (NAC, residues 61-95), which forms the fibril core; the familial Parkinson’s disease mutations H50Q and A53T accelerate aggregation while A30P and G51D slow it compared to the wild type (WT)^[24]^. Under the default decoder setting, mean Rg was nearly identical across the WT, H50Q, and A53T (38.51, 38.31, 38.42 Å), yet the high-NAC-exposure conformer fraction increased from 0.227 (WT) to 0.283 (H50Q) and 0.297 (A53T), with mean NAC shielding decreasing from 0.875 to 0.847 (H50Q) and 0.833 (A53T). Conversely, A30P and G51D decreased the high-NAC-exposure conformer fraction to 0.217 and 0.213 and increased mean NAC shielding to 0.995 and 1.011, respectively **(Supplementary Table S5)**. Residue-wise decomposition showed reduced NAC contacts in the accelerating mutants and the opposite for A30P and G51D, with the dominant redistribution in the proximal C-terminal region following the NAC (Figure 5e) — consistent with experimental links between C-terminal interactions and NAC exposure^[24]^. Notably, all four mutation sites lie outside the NAC region, indicating that sequence changes can alter NAC-associated contact organization without appreciably changing global compactness.

Together, these benchmarks show that MANTA-derived ensembles contain regional conformational information that is not reducible to global Rg. Decoder-driven expansion changes motif shielding in tau and Aβ, whereas sequence variants in α-synuclein redistribute local exposure populations at nearly unchanged global dimensions. These quantities should be interpreted as contact-defined ensemble descriptors rather than direct predictions of aggregation kinetics, mutation mechanisms, or disease phenotypes.

### 2.7. All-atom Reconstruction Enables Comparison with Experimental Observables

We next tested whether MANTA Cα ensembles could be reconstructed at all-atom resolution while preserving ensemble-scale properties. Cα-restrained rebuilding and energy minimization on IDPFold benchmark systems yielded all-atom ensembles that preserved the original ensemble-averaged Rg values (R^2^ = 0.9916, mean absolute difference 0.451 Å; **Figure 6a**). Structural relaxation remained modest overall (mean Cα-RMSD 1.834 Å; Figure 6b and **Supplementary Table S6**), indicating that restrained reconstruction retains the global organization of the input ensembles while providing atomistic coordinates for downstream observable calculations.

**Figure 6.**
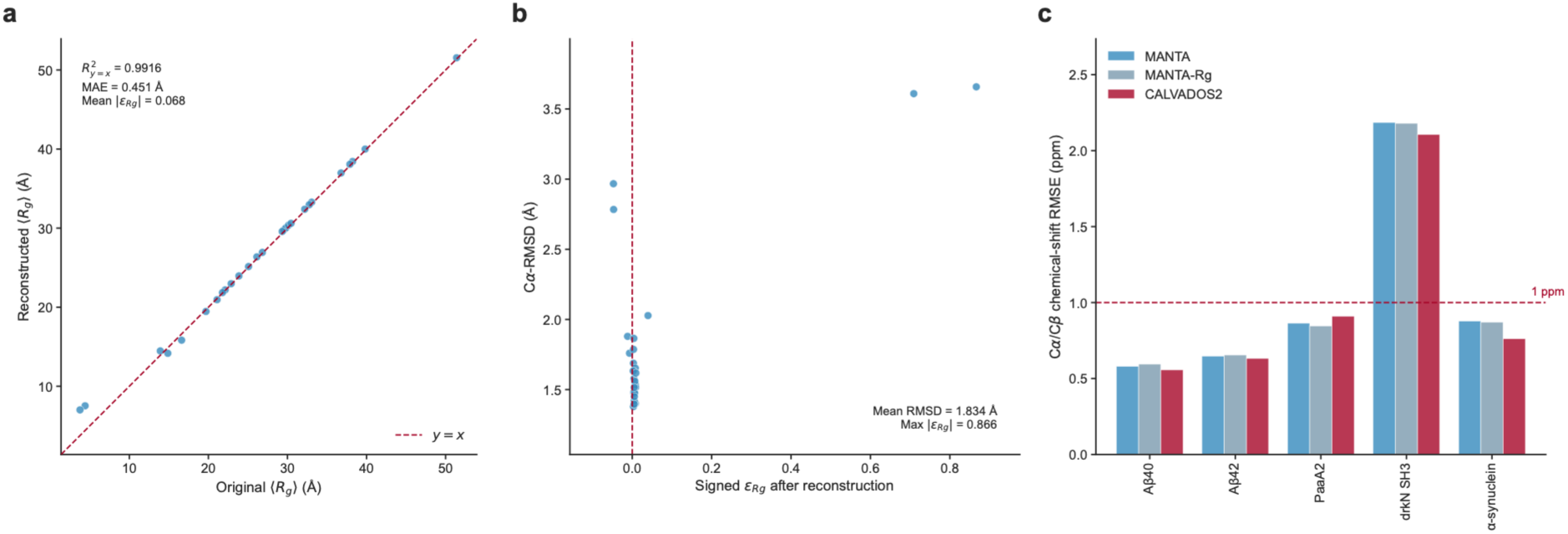
Cα-restrained all-atom reconstruction preserves ensemble-scale compactness and enables auxiliary observable comparison. **a.** Ensemble-averaged Rg values before and after all-atom reconstruction. **b.** Cα-RMSD after all-atom reconstruction plotted against the signed Rg change relative to the generated Cα-ensemble. **c.** Sequence-exact BMRB chemical-shift comparison after reconstruction. Bars show combined Cα/Cβ RMSE between SHIFTX2-predicted and experimental chemical shifts. MANTA-Rg denotes MANTA ensembles conditioned on reported IDPFold benchmark Rg values. The dashed line marks 1 ppm.

The reconstructed all-atom ensembles were then compared against SAXS and NMR data. Across five SAXS systems, Histatin5, PaaA2, Sic1, α-synuclein, and drkN SH3, default MANTA ensembles showed log-RMSE values of 0.052–0.265, and Rg-conditioning lowered them for two systems, most prominently Histatin5 (0.164 to 0.040) **(Supplementary Table S7)**. CALVADOS2, a CG model parameterized using experimental SAXS and Rg data, yielded lower log-RMSE for four systems, whereas MANTA yielded lower error for drkN SH3. Because MANTA ensembles were generated without SAXS fitting, reweighting, or refinement, these comparisons serve as out-of-distribution consistency checks, not like-for-like benchmarks.

For five NMR systems, Aβ40, Aβ42, PaaA2, drkN SH3, and α-synuclein, MANTA yielded sub-ppm pooled Cα/Cβ chemical shift RMSE (Figure 6c, **Supplementary Table S8**) for four systems (0.58–0.88 ppm), while drkN SH3 showed a higher error of approximately 2.2 ppm. CALVADOS2 showed a similarly elevated value for drkN SH3 (2.11 ppm), suggesting that this discrepancy is not specific to MANTA. CheSPI^[25]^-derived secondary-structure populations showed comparable agreement with Biological Magnetic Resonance Bank (BMRB)-derived profiles for the two methods, with system-dependent differences **(Supplementary Table S9)**.

Taken together, these results show that Cα-restrained reconstruction largely preserves MANTA ensemble-scale geometry and enables comparison with experimental observables. These analyses support the use of MANTA ensembles as templates for downstream atomistic reconstruction and observable calculation, rather than constituting coordinate-level experimental validation.

## 3. Discussion

Our results identify a specific limitation in sequence-based prediction of IDP ensembles. Sequence-derived representations captured substantial information about global compactness and inter-residue geometry. However, prediction accuracy decreased with increasing sequence separation. This suggests that long-range organization remains the dominant source of uncertainty. By separating geometric inference from conformational realization, MANTA enables the independent testing of these contributions. When reference-derived geometric priors were supplied to the same decoder, the median absolute relative Rg error decreased from 8.78% to 1.89%, accompanied by improvements in pairwise-distance, distributional, and contact-density agreement. The findings suggest that the majority of residual error in sequence-only generation stems from imperfect geometric priors rather than from coordinate realization.

This separation also reveals that ensemble organization cannot be reduced to a single compactness variable. The decoder controls altered mean chain dimensions and conformational breadth along partially separable axes, while matched-Rg controls demonstrated that changes in internal scaling were not explained by post hoc global rescaling alone. More importantly, local ensemble organization could vary independently of global size. This is evidenced by decoder-dependent changes in motif shielding, as shown by tau and Aβ. In addition, α-synuclein variants redistributed NAC-associated conformational populations despite nearly unchanged mean Rg. The direction of these local shifts was consistent with reported variant-dependent differences in aggregation behavior^[24]^, but these quantities remain contact-defined ensemble descriptors rather than direct predictions of aggregation kinetics or disease phenotypes.

MANTA therefore provides a framework for distinguishing global compactness, conformational heterogeneity, and local sequence-dependent organization as related but non-equivalent properties of IDP ensembles. The graph-based decoder enables rapid generation, producing 300 Cα-level conformers in a median of 17.14 s per system on the tested hardware. However, it should be noted that such runtimes are not directly comparable across methods due to variations in hardware, resolution, and implementation.

The Cα representation also defines the scope of the method. MANTA does not directly model side-chain packing, backbone torsional preferences, or atomic interactions. Cα-restrained all-atom reconstruction instead provides a template-completion step that preserves ensemble-scale compactness and enables calculation of auxiliary SAXS and NMR observables, rather than an independent atomistic equilibrium ensemble. The resulting comparisons support downstream compatibility but should not be interpreted as coordinate-level experimental validation. Physics-based CG approaches such as CALVADOS are therefore complementary, emphasizing physically parameterized sampling. Conversely, MANTA emphasizes sequence-derived geometry, rapid generation, and decoder-level controllability.

The decoder controls are currently effective parameters rather than physical observables. Changes in *p_Rg_* and ensemble width are not yet mapped to solvent conditions, ionic strength, or temperature. Furthermore, the external benchmark places significant reliance on Rg, which provides information regarding global chain dimensions rather than complete conformational distributions. More extensive validation against SAXS, additional NMR observables, FRET distances, mutation-dependent compaction, and environmental perturbations will be needed to establish condition-specific predictive accuracy.

In summary, separating geometric inference from conformational realization reveals long-range sequence-derived geometry as a major unresolved component of IDP ensemble prediction. MANTA uses this separation to provide controllable access to global dimensions, ensemble heterogeneity, and local conformational organization, offering a framework for examining how sequence variation and geometric uncertainty reshape disordered-protein conformational landscapes.

## 4. Method

### 4.1. Dataset Preparation

The training, validation, and test datasets were constructed from the PED^[13]^ (accessed April 25, 2026), which provides conformational ensembles for IDPs and IDRs, containing both experimentally determined and computationally modeled structures. The downloaded PDB-format ensemble files were used as the starting point. From each PED entry, the following data were extracted: the amino acid sequence, the number of chains, the number of conformations, the ligand information, and the file metadata. Single protein chain ensembles lacking a ligand were retained to concentrate on single-chain IDP/IDR ensembles and to circumvent confounding effects from intermolecular contacts. The PED files exhibiting identical amino acid sequences were merged into a unified non-redundant entry. For each unique sequence, source-file metadata were retained, and all conformational models were combined into a unified multi-model single PDB file to preserve all available conformations while removing exact sequence redundancy. The merged dataset was subsequently filtered to ensure compatibility with feature extraction, target generation, and model training. Entries that lacked essential information, such as sequence or ensemble model count, were discarded. To avoid the issue of excessive memory and preprocessing cost, the entries comprising more than 10,000 conformational models were excluded. Furthermore, given that ESM-2^[11]^ has an input length limit, sequences exceeding 1022 residues were removed. Sequences containing non-standard amino acid characters were also excluded.

To prevent data leakage to the external benchmark, the IDP entries exhibiting exact matches, overlapping constructs, or entries with at least 90% sequence identity and at least 90% benchmark-sequence coverage to any of the IDPFold benchmark sequences were excluded. This filtering was implemented using MMseqs2^[14]^ easy-search. The benchmark set was excluded from all training, validation, and calibration procedures. The final dataset comprised 6,755 entries from an initial set of 7,404 raw records, with a retention rate of 91.2%.

To minimize sequence leakage between the training, validation, and internal test sets, homology-aware splitting was employed based on sequence similarity. The pairwise sequence similarities between curated non-redundant sequences were computed using MMseqs2 easy-search. Pairs of sequences were designated as ‘related’ if they exhibited a minimum of 30% sequence identity and a minimum coverage threshold of 0.5. These similarity relationships were then used to construct an undirected graph, where nodes represent entries, and edges represent homologous relationships. The connected components of the graph were assigned to the training, validation, and internal test sets with a ratio of approximately 8:1:1, ensuring that homologous sequences were not split across different subsets. As a result, the homology-aware split comprised 5,404 training entries, 676 validation entries, and 675 internal test entries with median sequence lengths of 296.5, 267.5, and 255.0 residues, respectively. The median ensemble Rg values corresponding to the samples were 34.80 Å, 34.78 Å, and 33.70 Å **(Supplementary Table S10)**. The validation set was utilized for the selection of encoder checkpoints and the calibration of decoder protocols. Conversely, the internal test set was allocated for the final evaluation of sequence-derived geometric prediction and ensemble decoding performance.

The IDPFold benchmark^[9]^, comprising 27 experimentally characterized IDPs/IDRs with reported Rg values, was used as an external evaluation set. Protein names, sequences, and experimental Rg values were obtained from the Supporting Information of Zhu et al.^[9]^ Five proteins each were selected for SAXS and NMR chemical shift benchmarks. For each benchmark sequence, 300 conformers were generated in MANTA’s default sequence-only mode (*p_Rg_*=1.00) using validation-selected decoder parameters and sequence-derived geometric priors alone. Experimental Rg values were reserved exclusively for the purpose of evaluation, not for training, validation, decoder calibration, Rg scaling, or *p_Rg_* selection. A separate Rg-conditioned analysis supplied the reported experimental Rg to the decoder to assess coordinate realization under known global compactness. This target-conditioned mode was analyzed separately and was not used to select, tune, or replace the default benchmark setting.

For comparison, CALVADOS2^[5]^ ensembles were generated using the single-chain IDR workflow. Each protein was simulated as a single chain at 293.15 K, an ionic strength of 0.19 M, and pH 7.5, with both termini charged. Simulations were performed in a 50-nm cubic simulation box without periodic boundary conditions using the CUDA platform. Each trajectory was propagated for 2.1 × 10⁶ integration steps with a friction coefficient of 0.01 ps⁻¹, and coordinates were recorded every 7,000 steps, yielding 300 conformations per system.

### 4.2. Geometric Supervision Targets Derived from IDP Ensemble

The supervision targets were derived from conformational ensemble structures. Cα coordinates were extracted from all conformers of each ensemble, and for each residue pair, four geometric targets were stored: a 32-bin distance probability distribution, an expected distance, a distance-derived affinity score, and an entropy-based confidence score. Pairwise Cα distance matrices were discretized into 32 bins using a hybrid binning scheme – linearly spaced bins (0.615 Å interval) up to 8.0 Å and geometrically spaced bins for distances up to 120.0 Å – preserving local resolution while covering the broad long-range variation of IDP ensembles.

The distance-derived affinity score was computed using a Gaussian distance kernel (σ=2.0 Å), and the confidence score was computed from the normalized entropy of the distance distribution, so that sharply peaked distributions yielded higher confidence and diffuse ones lower. Local chain geometry was taken from ensemble-averaged *i* and *i+2* Cα distances, and global compactness from the ensemble-averaged Rg. The resulting precomputed files contained distance probability tensors, expected distance maps, local chain geometry, entropy-derived confidence maps, residue metadata, and ensemble-level Rg values.

### 4.3. Sequence Feature Extraction from a Protein Language Model

For a given input sequence, residue-level and pairwise representations are extracted using the pre-trained ESM-2 model (esm2_t33_650M_UR50D). For a sequence of length *L*, the residue-wise embeddings with a size of *L×1280* are obtained from the final hidden layer. The attention maps from the final four ESM-2 layers are also extracted. Given that each layer contains 20 attention heads, the resulting pairwise attention tensor is a size of *L×L×80*. Special beginning and ending tokens are removed from both embedding and attention outputs. The resulting ESM-2 features are reused during encoder training and evaluation.

### 4.4. Architecture of the Encoder Model

Following the extraction of ESM-2 residue embeddings from a given input sequence of length L, the embeddings are initially projected into a hidden representation comprising 128 dimensions. To construct pairwise features, the projected embeddings of residues *i* and *j*, denoted *h_i_* and *h_j_*, are expanded over all residue pairs and concatenated. Concurrently, the ESM-2 attention features for each residue pair are projected into the same hidden dimension, yielding *a_ij_*. The initial pairwise representation is subsequently obtained by concatenating the two projected residue embeddings and the projected attention features, followed by a linear fusion layer **(Equation 1)**:

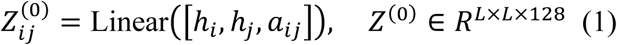

The fused pairwise representation is refined utilizing a stack consisting of three triangle multiplicative update blocks^[26]^. These blocks are designed to update residue-pair representations by aggregating information through pairwise relationships mediated by intermediate residues. This process enables the capture of higher-order pairwise relationships relevant to geometric consistency. After the refinement stack, the pairwise representation is symmetrized as **Equation 2**. This ensures that the predicted residue-pair features are invariant to the order of residue pairs.

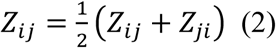

The refined pairwise representation is processed by four task-specific prediction heads. A distogram head predicts a 32-bin inter-residue distance distribution using a linear layer followed by log-softmax, while a distance regression head predicts the corresponding expected distance map using a so ftplus activation. A local separation head uses the off-diagonal pair representations *Z_i,i+2_* to estimate Cα distances between *i* and *i+2* residues. In parallel, a global Rg head applies adaptive average pooling and regression to estimate sequence-level compactness. For a given batch size *B* and sequence length *L*, the model produces the outputs as in **Equation 3**.

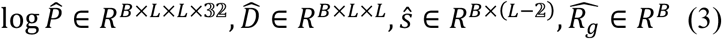

The model was trained end-to-end using a weighted multi-task objective comprising a KL-divergence loss for the predicted distance distribution and RMSE losses for the expected distance map, local *i, i+2* distances, and global Rg **(Equation 4)**.

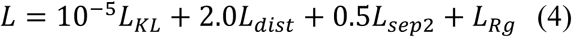

Here, *L_KL_* was computed between the target distance probability distribution and the predicted log-probabilities. *L_dist_*, *L_sep2_*, and *L_Rg_* denote RMSE losses for the expected distance map, local *i* and *i+2* distances, and global Rg, respectively. The distogram target was trained through the KL-divergence term, but no separate confidence-prediction head was trained.

The encoder was trained for up to 100 epochs using Adam^[27]^ with a learning rate of 10^-4^. The batch size was set to 2, with 4-step gradient accumulation, yielding an effective batch size of 8. The checkpoint exhibiting the lowest validation loss was selected for downstream ensemble generation. The training and validation metrics, including task-specific losses, distance MAE, Rg MAE, and Pearson correlation for Rg, were recorded at each epoch. The global Rg head was trained as an auxiliary compactness-prediction task but was not used to define the default decoder compactness target, which was instead derived from the predicted distance map.

### 4.5. Decoder Target Construction and Graph Parameterization

The mathematical decoder converts sequence-derived geometric outputs into Cα-level conformational ensembles. For an IDP sequence of length *L*, the decoder employs the predicted expected distance map, a distogram-derived pair confidence map, residue metadata, and an Rg-based compactness target. The pair confidence score *c_ij_* is computed as the maximum predicted probability across the distance distogram bins for residue pair *i* and *j*.

In the default sequence-only setting, the compactness is estimated from the predicted expected distance map using predicted pairwise distances as in **Equation 5**, rather than taken directly from the global Rg head.

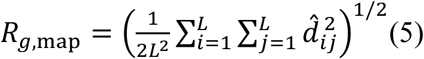

*d*^^^*_i_*_j_ is the predicted expected Cα distance between residues *i* and *j*. The global Rg head was trained as an auxiliary compactness-prediction task, whereas *R_g,map_* was used as the default sequence-only decoder compactness estimate.

The Rg-scale proxy, *p_Rg_*, functions as a decoder control variable for the mean target Rg, not a direct measurement of ensemble compactness. In the default sequence-only setting, the mean target Rg was obtained by multiplying *R_g,map_* by the validation-selected mean-scale factor, *s_mean_* and *p_Rg_* **(Equation 6)**.

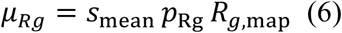

In the context of sequence-only mode, the parameter *p_Rg_* is set to 1.00, unless a specific value of the parameter is designated. When values are less than 1.00, compacting decoder settings are imposed, whereas when values are greater than 1.00, expanding decoder settings are imposed. The frame-level Rg sampling standard deviation, σ_Rg_, is fixed at 30% of the *R_g,map_* by default and is varied through the Rg width scale for controllability analyses. Consequently, changing *p_Rg_* results in a shift of the mean target size without directly broadening the Rg sampling distribution. For each conformer *m*, a frame-specific Rg value was sampled as in **Equation 7**, and clipped to remain within a geometrically plausible range.

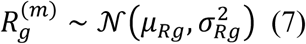

In instances when an explicit Rg is provided for target-conditioned generation, it is utilized as the mean target Rg for the decoder. During the benchmark study, the explicit Rg-conditioned mode was analyzed separately from the default sequence-only setting. This was due to the fact that the supplied target can contain experimental information.

For each conformer, residue-pair distance targets are sampled from the predicted expected distance map with confidence-dependent variability. Larger perturbations are applied to residue pairs with lower-confidence values, while high-confidence residue pairs are kept closer to the predicted expected distances. The perturbation uses spatially correlated Gaussian noise rather than independent per-pair noise, producing smoother conformational variation along the chain. Short-range residue pairs with high confidence scores serve as anchor constraints with stronger influence during coordinate realization. Conversely, the residue pairs with low confidence or long-range are treated more flexibly and guided by an ideal IDP-like polymer scaffold. For these flexible pairs, target distances follow Flory-type sequence-separation scaling^[18]^ set to the sampled frame-specific Rg, with optimization weights kept as low as possible. This preserves confident local and medium-range geometry while retaining flexible long-range disorder. Backbone connectivity is enforced separately. The *i, i+1* and *i, i+2* Cα distances are constrained using residue-dependent parameters from Martini-like CG bonded interactions^[28]^, ensuring chain continuity during distance-geometry reconstruction.

The sampled target distances and confidence scores are converted into a weighted residue-pair graph, with each residue as a node and residue-pair constraints as weighted edges. Distance-dependent affinity and confidence-dependent strengthening are combined into the edge weights *w_ij_* **(Equation 8)**.

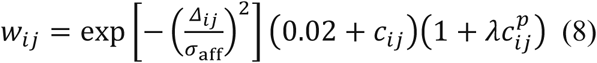

where *Δ_ij_* is the effective target distance, c_ij_ is the pairwise confidence score, σ_aff_=12.0 Å, λ controls confidence-dependent upweighting, and p is the confidence exponent. λ and p are decoder hyperparameters. To keep coordinate realization stable, residues with broadly uncertain distance profiles are down-weighted to prevent unstable hub-like constraints from dominating, and edge weights are capped so that no small set of constraints controls the graph. Residue pairs satisfying the anchor thresholds, *Δ_ij_ ≤* 20.0Å and *c_ij_ ≥* 0.15, act as anchor-like constraints and use the sampled target distance directly, whereas non-anchor or weakly constrained pairs instead use the IDP-like polymer scaffold distance with a weak sequence-separation-dependent weight. This weighted graph defines the Laplacian matrix used in graph-based distance-geometry realization, in which strong edges act as geometric anchors and weak edges give soft polymer-like guidance to disordered regions.

### 4.6. Graph-Based Coordinate Realization and Decoder Calibration

Initial Cα coordinates are generated using an isotropic fixed-step random walk and subsequently rescaled to correspond to the sampled frame-specific Rg. The coordinates are then determined under the constraints imposed by the weighted graph using a graph-based distance-geometry procedure. In the present implementation, a hierarchical SMACOF^[12]^-style weighted relaxation is employed, which iteratively minimizes the weighted stress between the target pairwise distances and the realized pairwise distances until convergence.

The process of structure generation comprises two stages. A high-confidence anchor graph is used to generate a stable initial configuration, followed by the full weighted graph for global relaxation, allowing both anchor constraints and weak polymer-scaffold terms to contribute to the final conformation. After the global relaxation, an Rg-guided local scaling procedure is initiated. Residue-wise flexibility is estimated from the predicted structural variability, and regions exhibiting greater variability are permitted to expand or contract more strongly. A short relaxation step is then used to restore consistency with the weighted distance constraints after local scaling, as the final step. After the coordinate realization, a chirality correction step is employed to circumvent mirror-image conformations. The net backbone chirality is estimated from consecutive Cα vectors, and mirrored conformations are corrected by reflecting the coordinates along the z-axis when necessary.

As relaxation and chirality correction may result in residual deviations from the targeted compactness, a final global Rg-matching step is applied as in **Equation 9**.

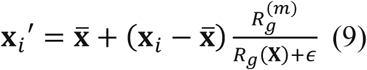

The term x-bar is defined as the center of the Cα coordinate, and *x_i_* and *x_i_’* are the Cα coordinates of residue i before and after rescaling. Rg(X) and Rg^(m)^ are the Rg values of the realized conformer before final scaling and the sampled target Rg for that conformer, respectively. ε is a small numerical stabilizer. This post-relaxation correction ensures that each realized conformer remains consistent with its intended compactness target while preserving the optimized relative geometry.

Decoder sampling parameters were selected on validation proteins and not updated using the internal test set or the external benchmark. Optuna^[15]^ selected four distance-sampling parameters (λ, p, hub weight scale, and sample mix) plus an additional mean-scale factor (s_mean_) for the default sequence-only Rg target. Each trial generated 20 conformers per validation protein, with the objective being the summed absolute relative errors in Rg and end-to-end distance. The selected parameters were λ=275.7387, p=6.9154, hub weight scale=0.3286, sample mix=0.9958, and s_mean_=1.1102. For default generation, the mean Rg target was therefore the distance-map-derived compactness estimate multiplied by 1.1102 and by the p_Rg_ (p_Rg_=1.00 defining the default sequence-only setting), and the frame-level Rg sampling width was fixed at 30% of that estimate. Encoder weights were not updated: calibration adjusted decoder sampling and mean size scaling parameters only, leaving the sequence-derived geometric predictions fixed.

### 4.7. All-atom Reconstruction and Application to SAXS Profile and NMR Chemical-Shift Analysis

All-atom models were generated from the Cα-level ensembles using PDBFixer and OpenMM^[29]^. For each conformer, PDBFixer was used to construct full-atom representations of each residue at pH 7.0. Missing residues inferred by the program were not added, restricting reconstruction to residues present in the input Cα trace. The resulting structures were subsequently minimized by OpenMM using the Amber14 force field^[30]^ and the OBC2^[31]^ implicit solvent model. Nonbonded interactions were treated with NoCutoff, and constraints were applied for the bonds attached to hydrogens. L-BFGS minimization^[32]^ was performed up to 500 iterations with a tolerance of 10 kJ·mol^-1^· nm^-1^. During minimization, harmonic positional restraints with a force constant of 10 kJ·mol^-1^· nm^-2^ were applied to Cα atoms, using the input Cα coordinates serving as reference positions. The minimized conformers were saved as multi-model PDB files. The effects of reconstruction were quantified using Cα-RMSD after Kabsch alignment^[33]^ and signed ε_Rg_ relative to the original Cα ensemble.

For SAXS comparison, theoretical scattering profiles of all-atom ensembles of Histatin5, PaaA2, Sic1, α-synuclein, and drkN SH3 were calculated using CRYSOL^[34]^. Experimental SAXS profiles for Histatin5 and α-synuclein were obtained from the Small Angle Scattering Biological Data Bank (SASBDB)^[35]^. For PaaA2, Sic1, and drkN SH3, CRYSOL profiles from PED were utilized for comparison. Ensemble-averaged scattering profiles were obtained by averaging the calculated profiles over all conformers. Experimental and predicted scattering profiles were compared over the common q range after interpolation onto the experimental q grid and normalization by the intensity at the lowest overlapping q value. Agreement was quantified using the log-RMSE between the normalized scattering profiles.

For the chemical-shift comparison benchmark, Cα/Cβ chemical shifts were predicted from reconstructed multi-model ensembles using SHIFTX2^[36]^ in NMR mode at pH 7.0 and 298 K. The five systems included in the study were Aβ40, Aβ42, PaaA2, drkN SH3, and α-synuclein, all of which exhibited exact sequence matches to BMRB^[37]^ entries. The predicted and experimental shifts were matched by residue index and atom name. RMSE values were calculated for Cα, Cβ, and pooled Cα/Cβ observations. The SHIFTX2-predicted chemical shifts were further interpreted by CheSPI^[25]^ to estimate residue-level helix, sheet, and coil population profiles. Sequence-exact BMRB chemical-shift entries were processed with the same CheSPI workflow to obtain reference profiles. A comparison was made between model-derived and BMRB-derived CheSPI profiles, with mean absolute differences calculated for each secondary structure population.

### 4.8. Evaluation Metrics

Generated Cα ensembles were evaluated using six metrics: global compactness, terminal distance, pairwise ensemble geometry, distance-distribution similarity, local structural validity, and controllability.

To measure global compactness, Rg was used. Rg of IDP with length L is calculated as **Equation 10** using Cα coordinates of residue i (x_i_):

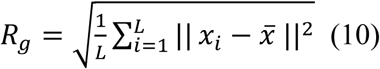

Comparison of the MANTA-generated ensemble Rg,gen with the experimental Rg,exp was conducted by calculating signed fractional Rg error, ε_Rg_, and its absolute value **(Equation 11)**.

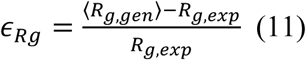

End-to-end distance was calculated as the Cα distance between the first and last residues. When reference ensembles were available, ensemble-averaged pairwise distance maps were compared using mean absolute error as in **Equation 12**.

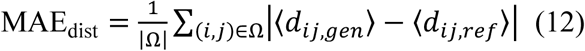

Ω denotes the evaluated residue-pair set. d_ij_ is a distance between residue i and j. By default, all pairs with *i<j* were included, whereas nonlocal analyses excluded short-range pairs with |*i*-*j*| < 3.

Pairwise distance distributions were compared using Kullback–Leibler divergence after discretization of experimental ensembles into the same 32 bins **(Equation 13)**.

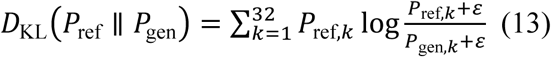

Here, ε was set to 10^-8^ to ensure numerical stability by preventing undefined logarithms and division by zero for zero-probability bins. KL divergence was calculated for each residue pair and averaged over evaluated pairs. Jensen–Shannon divergence was also used as a symmetric and bounded distributional metric **(Equation 14)**.

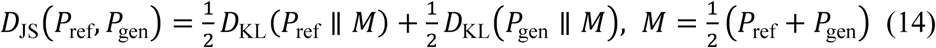

Contact density was defined as the fraction of nonlocal residue pairs (residues separated by more than 2) with Cα distance below 8.0 Å, averaged over conformers. In instances where reference ensembles were available, the contact density error was calculated as the absolute difference between the contact densities of generated and reference ensembles.

The local structural validity of Cα models was evaluated by using local chain-geometry criteria. A conformer was considered valid when Cα distances of all consecutive residue pairs fell within 2.8–4.8 Å and all *i, i+2* Cα distances fell within 4.0–8.5 Å. The consecutive Cα-distance criterion follows the same general principle as validity checks used in previous generative IDP ensemble models, which assess whether neighboring Cα distances remain physically plausible. We also included the *i, i+2* distance criterion as a local-geometry sanity check. The validity score was calculated as the fraction of conformers that satisfied both criteria.

The controllability of the decoder was assessed by systematically varying the decoder Rg-scale proxy p_Rg_, maintaining the frame-level Rg sampling width fixed. The alterations observed in the six metrics were measured across p_Rg_ values. It is important to note that values other than p_Rg_ of 1.00 were exclusively employed for the purpose of quantifying decoder controllability.

To distinguish decoder-induced changes in internal chain organization from trivial global size scaling, matched-Rg controls were constructed by isotropically rescaling default conformers about their Cα centroids to match the conformer-level Rg distribution of each Rg-scale-controlled ensemble, without rerunning the decoder. Apparent scaling exponents were then calculated using the same procedure as for the fully decoded ensembles. The quantification of sequence-separation scaling exponents was performed using the apparent Flory exponent, ν^[16]^, which describes the rate at which mean internal chain distance increases with increasing sequence separation. For each separation s, R(s) was defined as the mean Cα– Cα distance between residue pairs separated by s positions along the sequence. ν was estimated by fitting a slope in **Equation 15** over the specified sequence-separation range.

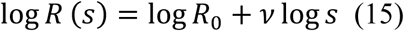

Short-range exponents were fitted over s from 1 to 10, whereas long-regime exponents were fitted over s ≥ 11 up to the maximum available sequence separation. As these fits were executed on finite generated ensembles, the interpretation of ν is interpreted as an apparent scaling descriptor rather than a universal polymer exponent was made.

For selected IDP cases, regional exposure was quantified using Cα-level contact descriptors. Cα contacts between a target region T and a set of shielding residues S were defined using an 8.0 Å distance cutoff, excluding residue pairs with sequence separation below 3. A binary contact-based exposure score, E_T_, was defined as one minus the fraction of residues in T forming at least one nonlocal Cα contact with S **(Equation 16)**.

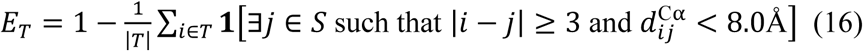

where 1 is the indicator function and *d*^C^**^α^** is the Cα-Cα distance. A contact-based shielding score S_T_ was defined as the number of nonlocal T-S contacts normalized by the number of residues in T **(Equation 17)**.

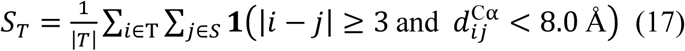

Higher E_T_ and lower S_T_ both indicate greater target-region exposure.

For the α-synuclein case study, the NAC region is residues 61–95, and S is all non-NAC residues. High-NAC-exposure conformers were those with S_T_ below the 25th percentile of the WT α-synuclein ensemble; this WT-referenced threshold was applied across WT, H50Q, A53T, A30P, and G51D ensembles to compare mutation-dependent shifts under a common definition. Target-region self-contact fraction, inter-region contact fraction, and mean inter-region Cα distance were also computed, with regions defined per system: N-terminal/NAC/C-terminal for synucleins, PHF6 and PHF6* for tau K18/K19, and CHC and C-terminal for Aβ40/Aβ42.

## Author Contributions

W.J. and W.S. conceived the study. W.J. developed the methodology and software, curated the data, performed the analyses, prepared the visualizations, and wrote the original manuscript. W.S. supervised and administered the project and reviewed and edited the manuscript. Both authors discussed the results and approved the final manuscript.

## Data Availability Statement

The curated training, validation, and test datasets, along with the external benchmark dataset used in this study, are publicly available on Zenodo at https://doi.org/10.5281/zenodo.23237426. The original protein ensemble data were obtained from the Protein Ensemble Database (PED; https://proteinensemble.org/). Experimental SAXS and NMR data used for evaluation were obtained from the Small Angle Scattering Biological Data Bank (SASBDB; https://www.sasbdb.org/) and the Biological Magnetic Resonance Bank (BMRB; https://bmrb.io/), respectively. The source code is available on GitHub at https://github.com/kucm-lsbi/MANTA and archived on Zenodo at https://zenodo.org/records/22122185.

Received: ((will be filled in by the editorial staff))

Revised: ((will be filled in by the editorial staff))

Published online: ((will be filled in by the editorial staff))

## Supporting Information

Supporting Information is available from the Wiley Online Library or from the author.

Intrinsically disordered proteins sample diverse conformations. MANTA uses protein language model features to infer geometric priors and a graph-Laplacian decoder to generate Cα ensembles. The modular design helps identify sources of error and provides partially separable controls over compactness and conformational breadth. The ensembles show sequence-dependent differences in contact-defined exposure of aggregation-associated motifs.

### ToC figure

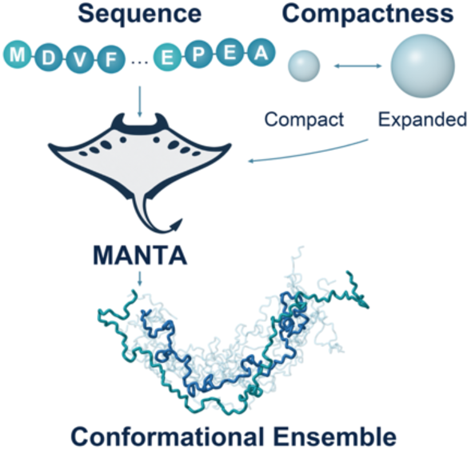

**Table S1.** Summary of MANTA performance under sequence-derived and reference-derived geometric inputs. Performance was evaluated on the homology-held-out test set. Predicted prior denotes sequence-only decoding from encoder-predicted geometric quantities, while reference prior denotes decoding from experimentally derived ensemble geometry using the same fixed decoder hyperparameters. Rg and end-to-end errors are reported as relative errors unless otherwise noted.

| **Metric** | **Predicted Prior** | **Reference Prior** |
|---|---|---|
| Rg signed relative error | 5.90% | 0.47% |
| Rg absolute relative error | 8.78% | 1.89% |
| End-to-end absolute relative error | 16.31% | 9.39% |
| Distance MAE all | 8.96 Å | 3.89 Å |
| Distance MAE nonlocal | 9.09 Å | 3.96 Å |
| JS divergence | 0.010 | 0.004 |
| Contact density error | 0.017 | 0.007 |
| Contact density PCC | 0.722 | 0.848 |

**Table S2.** External Rg benchmark of the default sequence-only operating point. Signed ε_Rg_ was calculated as the relative difference between the generated ensemble-averaged Rg and the experimental Rg. Values for previously published methods were taken from the IDPFold benchmark table. Because MANTA operates at Cα resolution, direct cross-method comparison was restricted to this resolution-compatible compactness metric. Experimental Rg values were used only for evaluation and were not provided to the decoder. Rg-scale proxy sweeps and explicit Rg-conditioned modes were analyzed separately and were not used to select, tune, or replace the default benchmark value. The methods that generate Cα-level ensemble are shown in bold.

| **Methods** | **Signed εRg** |
|---|---|
| **CALVADOS2** | 0.02 |
| **STARLING** | 0.06 |
| IDPFold | -0.06 |
| bAIes | -0.10 |
| a99SB-disp (7 systems) | 0.11 |
| AF-cluster | -0.12 |
| **idpGAN** | -0.12 |
| BioEmu | -0.18 |
| MD3*1µsESFF1+OPC3-B | -0.19 |
| AlphaFlow | -0.24 |
| **idpSAM** | -0.55 |
| MANTA | -0.11 |

**Table S3.**
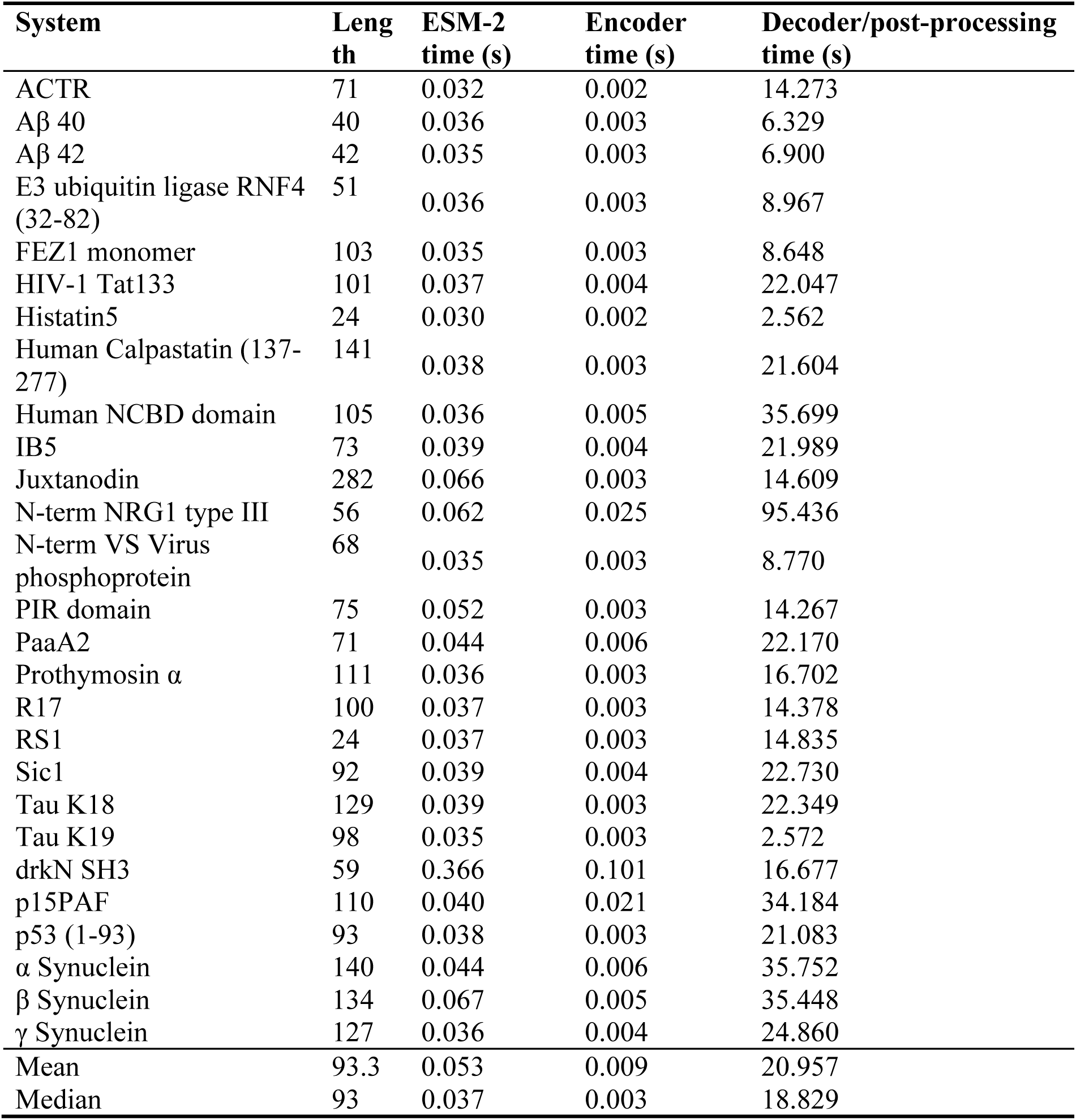
Per-system runtime characteristics for default sequence-to-ensemble generation. Runtime values were measured on the external 27-system benchmark using the default sequence-only setting, *p_Rg_* = 1.00, and 300 generated conformers per system. Timings were recorded after model loading. ESM-2 time denotes sequence feature extraction, encoder time denotes geometric-prior prediction, and decoder/post-processing time denotes graph-Laplacian ensemble decoding followed by multi-model PDB writing and rigid-core conformer alignment.

**Table S4.** System-level response to decoder Rg-scale modulation on the external IDPFold benchmark. Values show signed ε_Rg_ for ensembles generated at fixed Rg-scale proxy values *p_Rg_* = 0.70, 0.85, 1.00, 1.15, and 1.30. The *p_Rg_* = 1.00 column corresponds to the default sequence-only operating point reported in Table S2. Other columns quantify decoder controllability and were not used to select, tune, or replace the default benchmark setting.

| **System** | **Experimental Rg** | **Rg-scale Proxy** |  |  |  |  |
|---|---|---|---|---|---|---|
|  |  | 70% | 85% | 100% | 115% | 130% |
| ACTR | 25 | -0.358 | -0.204 | -0.115 | 0.014 | 0.152 |
| Aβ 40 | 12 | -0.099 | 0.059 | 0.239 | 0.449 | 0.649 |
| Aβ 42 | 12.4 | -0.076 | 0.121 | 0.340 | 0.503 | 0.684 |
| drkN SH3 | 16.7 | -0.402 | -0.263 | -0.167 | -0.036 | 0.080 |
| E3 ubiquitin ligase RNF4 (32-82) | 25.8 | -0.454 | -0.349 | -0.237 | -0.116 | -0.013 |
| FEZ1 monomer | 36 | -0.392 | -0.266 | -0.179 | -0.048 | 0.093 |
| Histatin5 | 13.8 | -0.775 | -0.728 | -0.681 | -0.614 | -0.579 |
| HIV-1 Tat133 | 33 | -0.356 | -0.209 | -0.097 | 0.042 | 0.204 |
| Human Calpastatin (137-237) | 39 | -0.381 | -0.253 | -0.152 | -0.020 | 0.121 |
| Human NCBD domain | 33 | -0.354 | -0.192 | -0.087 | 0.065 | 0.199 |
| IB5 | 27.9 | -0.399 | -0.261 | -0.145 | -0.014 | 0.082 |
| Juxtanodin | 55.9 | -0.336 | -0.219 | -0.081 | 0.066 | 0.216 |
| N-term NRG1 type III | 26.8 | -0.416 | -0.315 | -0.213 | -0.075 | 0.023 |
| N-term VS Virus phosphoprotein | 26 | -0.358 | -0.195 | -0.120 | 0.032 | 0.154 |
| p15PAF | 28.1 | -0.193 | -0.036 | 0.145 | 0.269 | 0.448 |
| p53 (1-93) | 28.7 | -0.270 | -0.090 | 0.061 | 0.198 | 0.335 |
| PaaA2 | 22.4 | -0.180 | -0.051 | 0.119 | 0.312 | 0.462 |
| PIR domain | 26.5 | -0.299 | -0.148 | 0.012 | 0.163 | 0.255 |
| Prothymosin α | 37.8 | -0.432 | -0.323 | -0.224 | -0.085 | 0.023 |
| R17 | 22.9 | -0.325 | -0.180 | -0.049 | 0.089 | 0.216 |
| RS1 | 12.62 | -0.786 | -0.743 | -0.702 | -0.654 | -0.609 |
| Sic1 | 34.7 | -0.452 | -0.336 | -0.247 | -0.098 | 0.020 |
| Tau K18 | 38 | -0.246 | -0.091 | 0.048 | 0.213 | 0.406 |
| Tau K19 | 35 | -0.312 | -0.229 | -0.064 | 0.076 | 0.217 |
| α Synuclein | 33 | -0.215 | -0.048 | 0.159 | 0.328 | 0.507 |
| β Synuclein | 49 | -0.489 | -0.382 | -0.249 | -0.164 | -0.026 |
| γ Synuclein | 61 | -0.556 | -0.450 | -0.379 | -0.276 | -0.196 |

**Table S5.** Global and NAC-exposure metrics for WT, H50Q, A53T, G51D, and A30P α-synuclein ensembles.

| **Variant** | **Mean Rg (Å)** | **Mean NAC shielding** | **High-NAC-exposure fraction** | **NAC-exposed extended fraction** | **NAC-exposed terminally released fraction** |
|---|---|---|---|---|---|
| WT | 38.51 | 0.875 | 0.227 | 0.093 | 0.090 |
| H50Q | 38.31 | 0.847 | 0.283 | 0.157 | 0.137 |
| A53T | 38.42 | 0.833 | 0.297 | 0.127 | 0.123 |
| G51D | 36.78 | 1.011 | 0.213 | 0.100 | 0.100 |
| A30P | 37.19 | 0.995 | 0.217 | 0.110 | 0.090 |

**Table S6.** System-wise comparison between original Cα ensembles and Cα-restrained all-atom reconstructed ensembles. Cα-RMSD measures the coordinate deviation between the original and reconstructed Cα traces. Signed ε_Rg_ (vs Original) denotes the relative change in ensemble-averaged Rg after all-atom reconstruction using the original Cα ensemble as the reference. Signed ε_Rg_ (vs Experimental) denotes the relative error of the reconstructed ensemble-averaged Rg against experimental Rg values.

| **System** | **Cα-RMSD(Å)** | **Signed εRg(vs Original)** | **Signed εRg(vs Experimental)** |
|---|---|---|---|
| ACTR | 1.688 | 0.003 | -0.113 |
| Aβ 40 | 2.784 | -0.047 | 0.181 |
| Aβ 42 | 2.967 | -0.047 | 0.276 |
| drkN SH3 | 2.028 | 0.040 | -0.134 |
| E3 ubiquitin ligase RNF4 32-82 | 1.880 | -0.012 | -0.246 |
| FEZ1 monomer | 1.401 | 0.008 | -0.172 |
| Histatin5 | 3.609 | 0.709 | -0.454 |
| HIV-1 Tat133 | 1.534 | 0.008 | -0.090 |
| Human Calpastatin 137-237 | 1.403 | 0.007 | -0.147 |
| Human NCBD domain | 1.653 | 0.008 | -0.080 |
| IB5 | 1.865 | 0.004 | -0.141 |
| Juxtanodin | 1.379 | 0.003 | -0.078 |
| N-term NRG1 type III | 1.759 | -0.007 | -0.218 |
| N-term VS Virus phosphoprotein | 1.577 | 0.004 | -0.116 |
| p15PAF | 1.559 | 0.007 | 0.153 |
| p53 1-93 | 1.419 | 0.005 | 0.067 |
| PaaA2 | 1.633 | 0.003 | 0.122 |
| PIR domain | 1.478 | 0.004 | 0.016 |
| Prothymosin α | 1.515 | 0.008 | -0.218 |
| R17 | 1.787 | 0.003 | -0.046 |
| RS1 | 3.658 | 0.866 | -0.444 |
| Sic1 | 1.618 | 0.009 | -0.240 |
| Tau K18 | 1.482 | 0.004 | 0.053 |
| Tau K19 | 1.475 | 0.006 | -0.058 |
| α Synuclein | 1.515 | 0.005 | 0.165 |
| β Synuclein | 1.450 | 0.005 | -0.245 |
| γ Synuclein | 1.396 | 0.005 | -0.376 |

**Table S7.** SAXS-profile comparison for reconstructed ensembles. MANTA-Rg denotes MANTA ensembles conditioned on reported IDPFold benchmark Rg values. Histatin5 and α-synuclein were compared with experimental SAXS profiles, whereas PaaA2, Sic1, and drkN SH3 were compared with PED-associated CRYSOL reference profiles. These comparisons were post hoc and were not used for training, SAXS fitting, reweighting, or refinement.

| **System** | **Ref.** | **MANTA log-RMSE** | **MANTA-Rg log-RMSE** | **CALVADOS2 log-RMSE** |
|---|---|---|---|---|
| Histatin5 | SASDHF8 | 0.164 | 0.040 | 0.033 |
| PaaA2 | PED00013e001 | 0.053 | 0.059 | 0.042 |
| Sic1 | PED00159e002 | 0.101 | 0.090 | 0.011 |
| α-Synuclein | SASDQJ7 | 0.265 | 0.270 | 0.193 |
| drkN SH3 | PED00158e006 | 0.052 | 0.052 | 0.100 |

**Table S8.** Sequence-exact BMRB chemical-shift comparison after all-atom reconstruction. SHIFTX2-predicted Cα/Cβ shifts from reconstructed ensembles were compared with experimental BMRB values for five sequence-exact systems. MANTA-Rg denotes ensembles conditioned on reported IDPFold benchmark Rg values. N(Cα/Cβ) denotes matched Cα and Cβ observations. RMSD values are reported in ppm. This analysis was used as an auxiliary NMR-observable assessment, not as coordinate-level structure validation.

| **System** | **BMRB ID** | **N(Cα/Cβ)** | **Method** | **Cα RMSE (ppm)** | **Cβ RMSE (ppm)** | **Cα/Cβ RMSE (ppm)** |
|---|---|---|---|---|---|---|
| Aβ40 | 17796 | 40/34 | MANTA | 0.54 | 0.63 | 0.58 |
|  |  |  | MANTA-Rg | 0.57 | 0.63 | 0.59 |
|  |  |  | CALVADOS2 | 0.52 | 0.60 | 0.56 |
| Aβ42 | 17794 | 42/35 | MANTA | 0.50 | 0.79 | 0.65 |
|  |  |  | MANTA-Rg | 0.50 | 0.80 | 0.66 |
|  |  |  | CALVADOS2 | 0.50 | 0.76 | 0.63 |
| PaaA2 | 18841 | 71/71 | MANTA | 1.11 | 0.52 | 0.86 |
|  |  |  | MANTA-Rg | 1.09 | 0.50 | 0.85 |
|  |  |  | CALVADOS2 | 1.21 | 0.44 | 0.91 |
| drkN SH3 | 25501 | 56/53 | MANTA | 1.85 | 2.49 | 2.19 |
|  |  |  | MANTA-Rg | 1.84 | 2.49 | 2.18 |
|  |  |  | CALVADOS2 | 1.78 | 2.40 | 2.11 |
| α-synuclein | 6968 | 140/122 | MANTA | 1.05 | 0.64 | 0.88 |
|  |  |  | MANTA-Rg | 1.04 | 0.63 | 0.87 |
|  |  |  | CALVADOS2 | 0.87 | 0.62 | 0.76 |

**Table S9.** CheSPI-derived local secondary-structure population comparison against BMRB-derived references. SHIFTX2-predicted chemical shifts from reconstructed MANTA, MANTA-Rg and CALVADOS2 ensembles were converted to CheSPI secondary-structure population profiles and compared residue-wise with CheSPI profiles derived from sequence-exact BMRB chemical shifts. Values report mean absolute differences for helix, sheet and coil populations. Lower values indicate closer agreement with the BMRB-derived profile. MANTA-Rg denotes MANTA ensembles conditioned on the reported benchmark Rg values.

| **System** | **CheSPI population** | **MANTA** | **MANTA-Rg** | **CALVADOS2** |
|---|---|---|---|---|
| Aβ40 | Helix | 0.052 | 0.056 | 0.050 |
|  | Sheet | 0.057 | 0.060 | 0.056 |
|  | Coil | 0.078 | 0.085 | 0.074 |
| Aβ42 | Helix | 0.058 | 0.067 | 0.054 |
|  | Sheet | 0.059 | 0.065 | 0.057 |
|  | Coil | 0.083 | 0.094 | 0.079 |
| PaaA2 | Helix | 0.321 | 0.318 | 0.329 |
|  | Sheet | 0.084 | 0.083 | 0.068 |
|  | Coil | 0.291 | 0.288 | 0.315 |
| α-synuclein | Helix | 0.239 | 0.235 | 0.176 |
|  | Sheet | 0.088 | 0.088 | 0.100 |
|  | Coil | 0.286 | 0.282 | 0.237 |
| drkN SH3 | Helix | 0.083 | 0.084 | 0.080 |
|  | Sheet | 0.060 | 0.060 | 0.059 |
|  | Coil | 0.111 | 0.112 | 0.116 |

**Table S10.** Statistics of the final train, validation, and internal test splits.

| **Split** | **Sequences** | **Median sequence length** | **Length range** | **Median conformers per entry** | **Total conformers** | **Median ensemble Rg** |
|---|---|---|---|---|---|---|
| Train | 5,404 | 296.5 | 5–825 | 100 | 584,623 | 34.8 |
| Validation | 676 | 267.5 | 42–808 | 100 | 69,252 | 34.78 |
| Internal test | 675 | 255 | 54–657 | 100 | 68,700 | 33.7 |
| Total | 6,755 | — | 5–825 | — | 722,575 | — |

**Figure S1.**
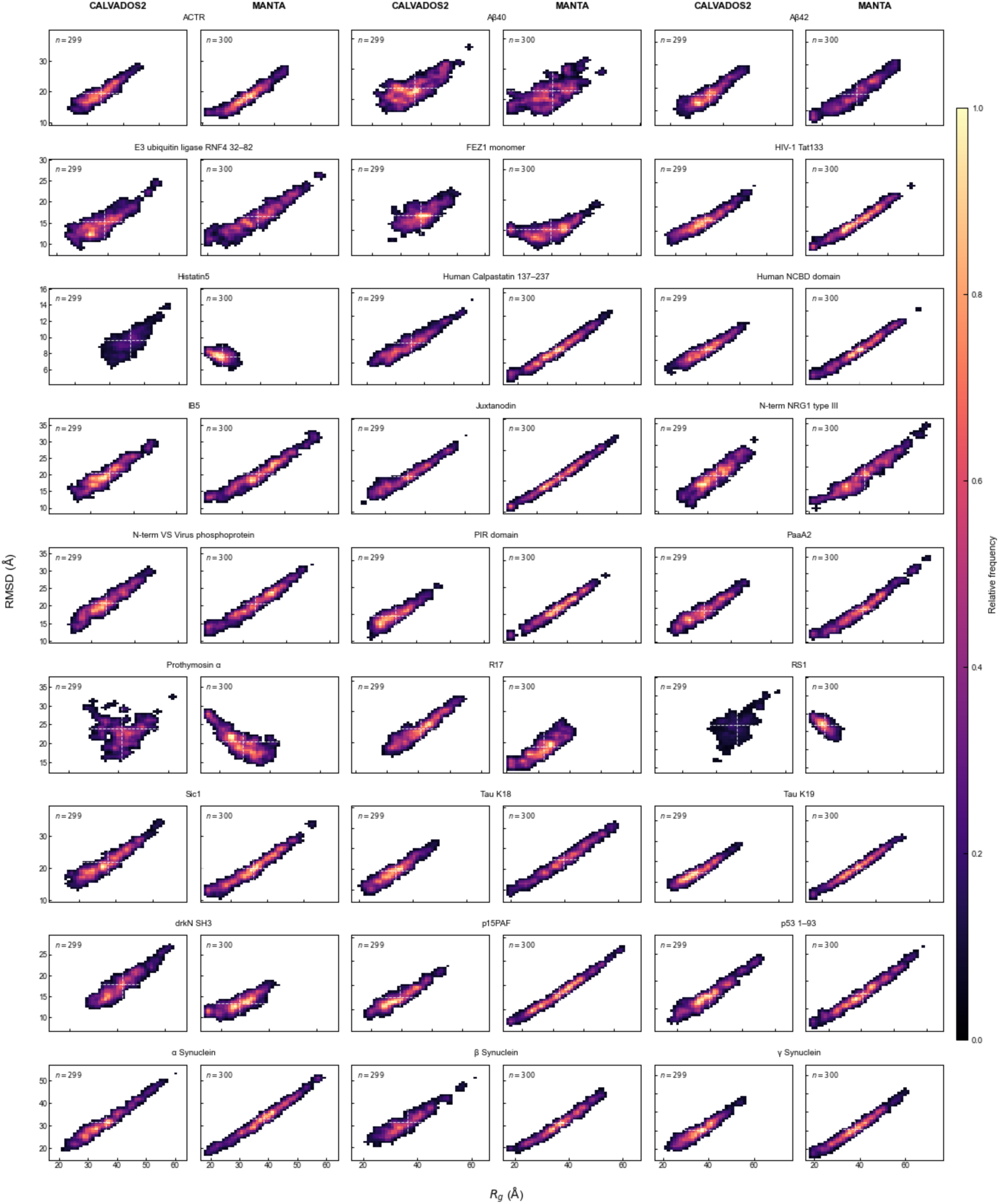
Rg–RMSD occupancy maps comparing MANTA and CALVADOS2 ensembles across the 27-system IDPFold benchmark. For each system, conformers were projected using radius of gyration and RMSD to the first CALVADOS2 model after optimal Cα superposition. The first CALVADOS2 model, used as the RMSD reference, was excluded from the density maps. Heat intensities indicate relative conformer frequency normalized within each system across the paired CALVADOS2 and MANTA maps. RMSD was used only as a visualization coordinate and not as a direct accuracy metric.

